# Independent history traces for distinct percepts derived from a single stimulus

**DOI:** 10.64898/2026.06.18.733184

**Authors:** Alessandro Toso, Mathew E. Diamond

## Abstract

Perception is more than a direct readout of current sensory input; it is also shaped by recent experience, a phenomenon known as serial dependence. While the effects of recent experience have been well characterized for single stimulus features, it remains unclear how perceptual history operates when multiple perceptual dimensions must be extracted from a single sensory event. Here, human participants reported their judgment of vibrotactile stimuli, with the relevant feature – either the intensity or duration - randomly cued on each trial. Within trials, intensity and duration interacted bidirectionally - longer stimuli were perceived as stronger and stronger stimuli as longer - indicating that both features are derived from a shared sensory representation. Across trials, however, serial dependence was feature-specific: perceived intensity was selectively attracted toward previously perceived intensities, and perceived duration toward previously perceived durations. The current trial judgment was influenced by perceptual estimates from multiple preceding trials, but only those in which the same feature was task-relevant. We formalize these findings with a computational model in which perceptual history is maintained in parallel memory traces, each selectively updated when the corresponding feature is task-relevant. The model captured both the psychometric structure of behavior and the feature-specific, multi-trial dependence of perceptual judgments, with traces that accumulate past percepts and bias current estimates along the corresponding feature dimension.Serial dependence in this task operates on perceptual representations downstream of a shared sensory encoding stage, and is maintained simultaneously in feature-specific, task-gated memory traces within higher-level cortical circuits.

## Introduction

Perceptual judgments are systematically shaped by recent experience. Responses on a given trial are biased by preceding stimuli across a wide range of tasks in both humans^1–7^ and animals^8–16^. These history effects can be either attractive or repulsive. In continuous estimation paradigms, perceived values are biased toward recently encountered stimuli (“history-dependent attraction”)^17–19^. This pattern is often interpreted as a form of temporal integration, whereby recent sensory inputs are weighted alongside current evidence when forming an estimate, effectively smoothing moment-to-moment variability in the sensory signal ^20,21^. By contrast, repulsive biases often arise in categorization tasks that require comparing sensory evidence to an internal decision criterion^3,11,16^. In such cases, recent stimuli attract the decision criterion toward those sensory values, causing subsequent judgments to be displaced away from the recent past (e.g., strong recent stimuli increase the likelihood of reporting the next stimulus as weaker). Thus, the sign and magnitude of history effects depend on how sensory information is read out at the decision stage – a property that is itself determined by task structure, rather than by the stimulus sequence alone.

Here, we focus on serial dependence in a continuous estimation task because this format provides a direct, graded readout of perceptual representations. Unlike categorical judgments, which can conflate perceptual and decision-level processes, continuous reports allow trial-by-trial quantification of biases induced by recent sensory history, enabling a more direct assessment of how past inputs influence current perceptual estimates. With the exception of a small number of studies^22,23^, serial dependence has been studied for single stimulus features (e.g., orientation, motion, numerosity, pitch, intensity, or duration) ^18,21,22,24–27^. Yet individual stimuli typically support multiple percepts. Vibrotactile stimuli, used in the present study, give rise to at least two dissociable perceptual dimensions - intensity and duration – both of which obey standard psychometric relations^14,28–31^. These percepts, are not independent: higher-intensity stimuli are perceived as longer, and longer stimuli as more intense^14,28–31^. Computational accounts reproduce these interactions by modeling percept formation as temporal accumulation of sensory drive with distinct integration time constants for each feature - shorter for intensity, longer for duration^28,30^.

At what representational level does perceptual history operate? When a single sensory event affords multiple perceptual attributes, does recent experience update a single unified history trace tied to the physical stimulus, or instead separate feature-specific traces? If the latter, do they track each feature whenever it is available in the stimulus, or only when that feature is explicitly estimated and reported?

We tested this idea with a *direct-estimation* task^32^ in which participants reported either the perceived intensity or perceived duration of vibrotactile stimuli, with the task-relevant feature randomly cued on each trial. This design decouples the history of the physical stimulus from the history of the reported perceptual estimate, allowing us to distinguish between a unitary stimulus-bound prior, feature-specific priors updated by stimulus availability, and feature-specific priors updated selectively according to the reported estimate.

## Results

### Reciprocal perceptual biases for intensity and duration

To quantify serial dependencies in perceived intensity and perceived duration, we used a direct estimation task (as in ^32^ ). The stimulus set comprised six durations (80–800 ms, linearly spaced) and six intensities (mean speed 9.6–67.2 mm/s, linearly spaced), yielding 36 feature combinations, all presented in each session (Figure 1A). On each trial, one duration/intensity combination was selected at random. Each trial began with a color-cue indicating the feature to be estimated – intensity (*I*; red) or duration (*T*; blue). After a 500-ms delay, a single vibrotactile stimulus, defined by *I* and *T*, was delivered to the left index fingertip. A response bar (slider) appeared 500 ms after stimulus offset (Figure 1A), and participants reported the magnitude of the cued percept by mouse-clicking a chosen position along the slider. To dissociate perceptual-history effects from potential motor carryover, slider orientation (vertical vs. horizontal) was randomized across trials, independently of the task cue. Scale normalization procedures are detailed in Methods.

**Figure 1.**
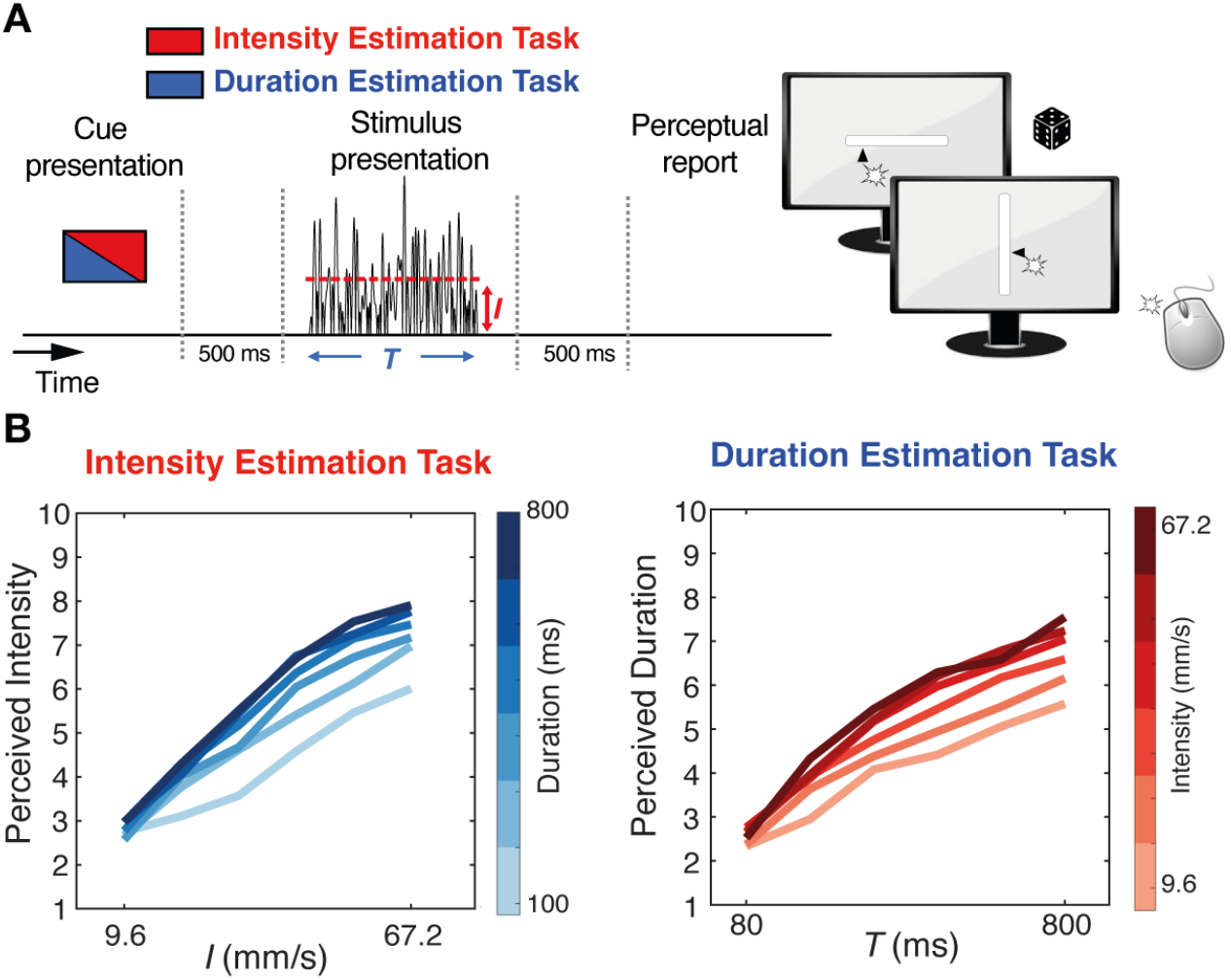
Interaction between intensity and duration. **(A)**Experimental setup: participants received a single vibrotactile stimulus and reported either its perceived duration or perceived intensity by clicking on a slider displayed on the computer monitor. A cue at trial onset indicated which feature had to be estimated. The orientation of the response slider (vertical or horizontal) was randomized on every trial. **(B)**Mean perceived intensity (left) and perceived duration (right) increased as a function of both the relevant (cued) feature and the irrelevant feature. Mean values are the grand average across subjects.

We first examined the reciprocal influence of the two stimulus features on perceptual judgments. The property cued for judgment is referred to as the “relevant” feature and the accompanying, uncued property as the “irrelevant” feature. As shown in Figure 1B-C, each feature biased judgments of the other: longer vibrations were judged as stronger (“longer feels stronger”), and higher-intensity vibrations were judged as lasting longer (“stronger feels longer”), replicating previously reported bidirectional perceptual coupling^32^. This cross-feature bias indicates that judgments draw on partially overlapping internal representations.

### Attraction of the current percept toward the previous percept

Overlap at the representational level (Figure 1) admits three alternative hypotheses about the origin of serial dependence, distinguished by the stage at which past information exerts its influence. First, history effects may arise at the level of *physical stimulus features*, such that the objective properties of prior stimuli directly bias subsequent judgments; under this account, any cross-feature transfer would be driven by the objective magnitude of intensity and duration. Second, history effects may instead arise within shared perceptual representations, in which case prior stimuli would bias the *perceived* attributes of subsequent stimuli; cross-feature transfer would then reflect the interacting perceptual representations of intensity and duration, rather than their physical values. Third, history effects may emerge downstream at the level of feature-specific readouts, such that serial dependence reflects how individual features are selectively accessed for judgment; in this case, serial dependencies should remain largely independent for intensity and duration, with little cross-feature transfer. To test these predictions, we focused first on the influence of the immediately preceding trial. Specifically, we asked how perceived intensity and duration on trial *n* depend on the intensity and duration of trial *n-1*, restricting the analysis to the condition in which *trials n-1* and *n* cued the same feature as relevant.

Figure 2A illustrates the case in which *n-1* and *n* were both intensity-estimation trials. Perceived intensity on trial *n* increased systematically with both the duration of the preceding stimulus, *T*_*n-1*_ (left panel), and its intensity, *I*_*n-1*_ (right panel)). Figure 2B summarizes these effects, defining the *n-1*-induced history bias as the slope relating *T*_*n-1*_ (blue-gray) and *I*_*n-1*_ (dark red) to perceived intensity on trial *n*. Both features exerted attractive influences, although the effect of intensity was substantially stronger (*p* = 0.0009, one-sided permutation test; see Methods).

**Figure 2.**
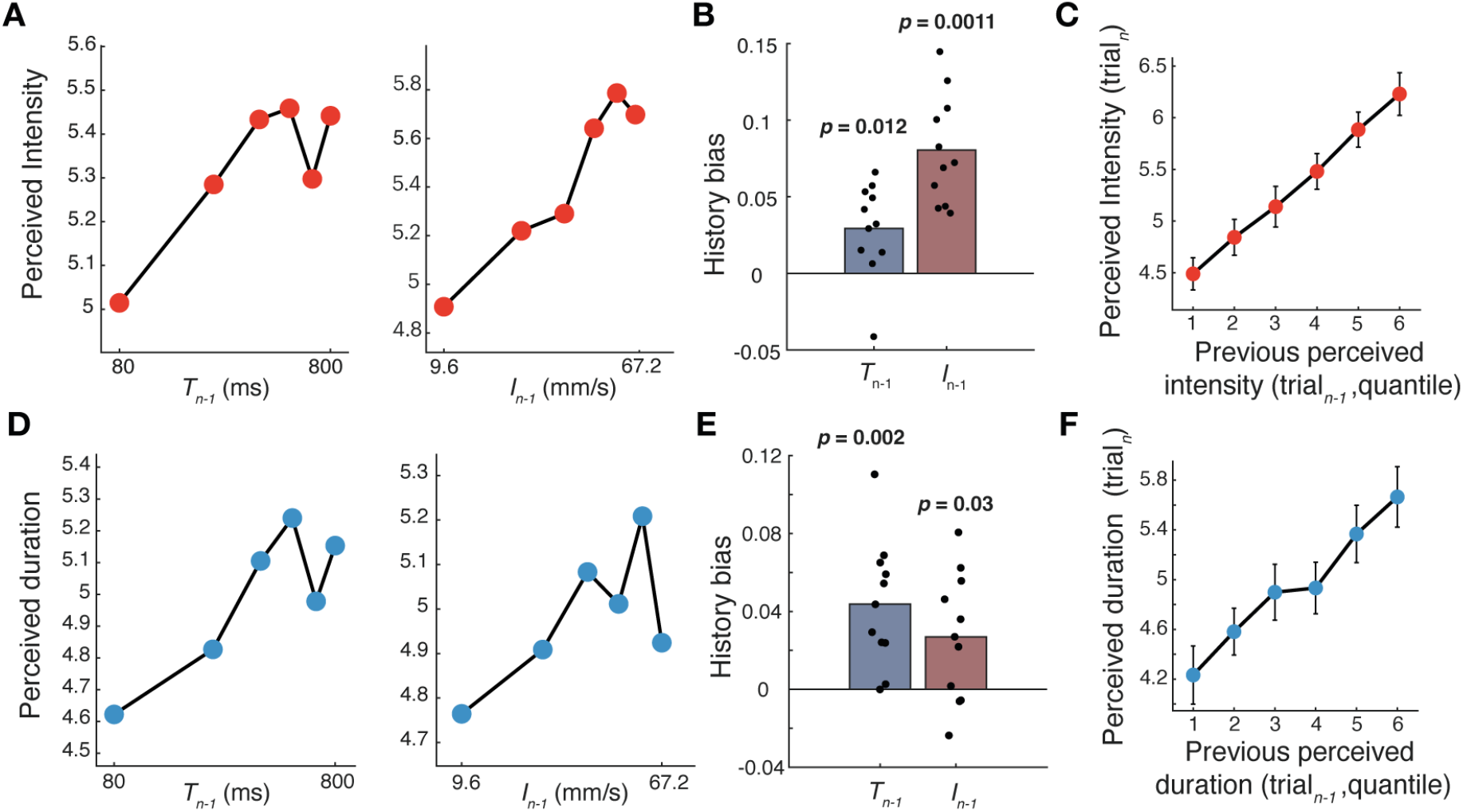
Serial dependencies when the cued feature is repeated on consecutive trials. **(A)**Mean perceived intensity on trial *n* plotted against the duration (left panel) and intensity (right panel) of stimulus *n-1*. **(B)**Duration-history bias (blue) and intensity-history bias (red) for all subjects; each point is one subject. *p*-values, permutation tests against zero. **(C)**Mean perceived intensity on trial *n* plotted against the perceived intensity on trial *n-1*. **(D)**Mean perceived duration on trial *n* plotted against the duration (left panel) and intensity (right panel) of stimulus *n-1*. **(E)**Duration-history bias (blue) and intensity-history bias (red) for all subjects; each point represents one subject. *p*-values, permutation tests against zero. **(F)**Mean perceived duration on trial *n* plotted against the perceived duration on trial *n-1*.

Thus, when intensity was estimated on consecutive trials, perceived intensity on trial *n* was more strongly attracted toward the relevant feature (intensity) of the preceding trial than toward the irrelevant feature (duration) (Figure 2B).

This raises the question, in light of the physical-versus perceptual-level accounts introduced above, of whether attraction tracks the physical properties of the preceding stimulus or the perceived magnitude extracted from it on trial *n-1*. Because the slider provides a direct readout of perceived magnitude, history effects can be expressed in perceptual units. Figure 2C shows that perceived intensity on trial *n* depends more strongly on perceived intensity on trial *n-1* than on the corresponding physical intensity (compare with right panel of Figure 2A), indicating that attraction is more robust in perceptual than in physical coordinates (*p* < 0.001, two-sided permutation test).

Figure 2D–F presents the analogous analysis for duration estimation. Perceived duration on trial *n* increased systematically with both preceding duration, *T*_*n-1*_ (left panel), and preceding intensity, *I*_*n-1*_ (right panel). As in the sequential intensity trials, these effects were asymmetric: preceding duration exerted a stronger influence than preceding intensity (*p* = 0.02, one-sided permutation test; Figure 2E). Consistent with the intensity analysis, perceived duration on trial *n* depended more strongly on perceived duration on trial *n-1* than on the corresponding physical measure in ms (Figure 2F vs. left panel Figure 2D, *p* < 0.001, two-sided permutation test). These results suggest that sequential attraction originates in the brain’s representation of the perceived experience, rather than in early, pre-conscious encoding of stimulus features. Supplementary Figures 2 and 3 confirm that all the effects seen in Figure 2 persisted independently of the orientation of the response slider across consecutive trials, indicating that sequential attraction cannot be attributed to a simple motor carryover.

To confirm that the influence of the preceding trial (*n-1*) is better explained by the percept extracted from stimulus *n-1* than the physical properties of *n-1*, we directly compared model fits across subjects. In sequences in which the cued feature was repeated, a model incorporating the previous perceptual report consistently explained more variance (higher *R*^*2*^) than a model based on the previous stimulus properties (Figure 3A). This result indicates that serial dependence is more strongly anchored to preceding perceptual representations than to the preceding physical features.

**Figure 3.**
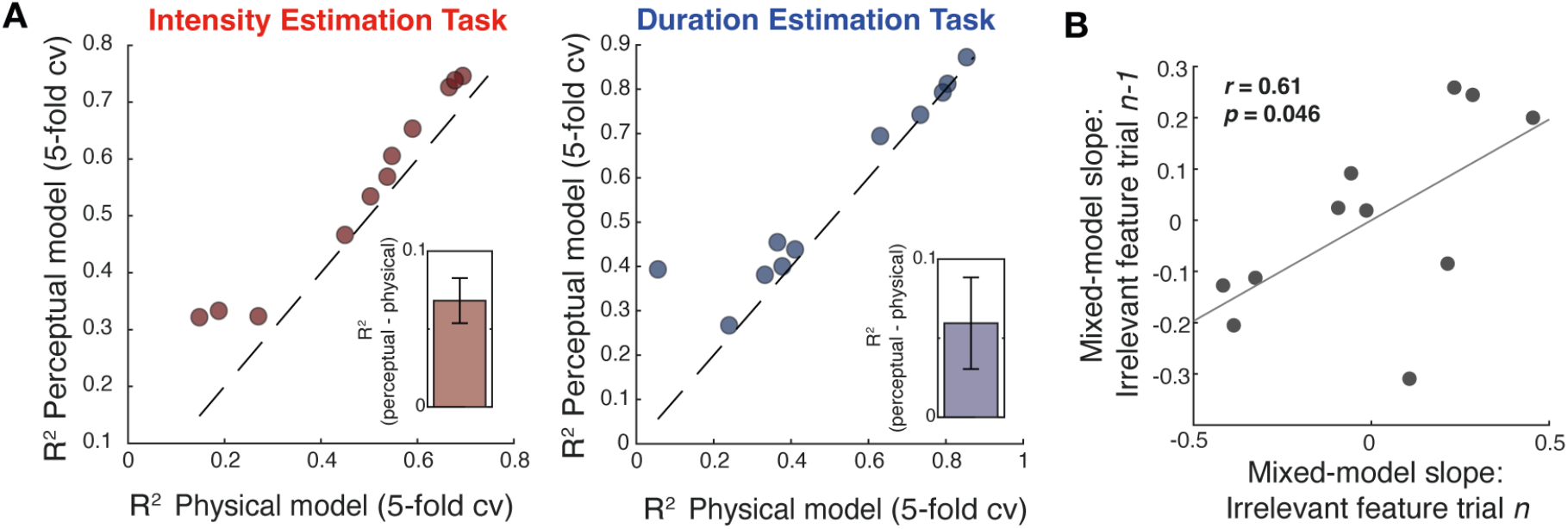
Influence of relevant and irrelevant features. **(A)**Cross-validated model comparison for the intensity task (left) and duration task (right). Each point represents one participant. The x-axis shows the variance explained (*R*^*2*^, 5-fold cross-validation) by a model incorporating previous physical stimulus features, whereas the y-axis shows the variance explained by a model incorporating previous perceptual reports. Points above the identity line indicate superior performance of the perceptual-history model. Insets depict the mean difference in *R*^*2*^ (perceptual - physical) across participants (error bars: ± SEM). **(B)**Correlation between subject-specific random slopes for the irrelevant feature on the current trial (trial *n*) and the previous trial (trial *n-1*).

If the influence of irrelevant features is not purely stochastic but reflects real differences in how observers weight sensory dimensions, then individuals who integrate these features more strongly should also exhibit stronger trial-to-trial carry-over. Using a linear mixed-effects model with random slopes for current and previous irrelevant features, we obtained subject-specific slope estimates and examined their relationship. A significant positive correlation emerged between the weights assigned to the irrelevant feature on the current trial and preceding trial (*r* = 0.61, *p* = 0.046; Figure 3B). Thus, individuals who integrate the irrelevant feature more strongly into their current percept also exhibit stronger carry-over of that feature to the subsequent trial. Together, the model comparison (Figure 3A) and the mixed-effects analysis (Figure 3B) indicate that serial dependence is more strongly tied to previous perceptual reports than to the physical properties of preceding stimuli, implicating perceptual representations as the relevant substrate of history effects.

### Parallel updating of independent history traces for intensity and duration perception

Figure 2 established that history effects are expressed in perceptual rather than physical coordinates, ruling out accounts based on the direct influence of physical stimulus features. We next asked whether these perceptual traces arise automatically from stimulus exposure or instead require the feature to be actively accessed for judgment. Under a perceptual-representation account, any encoded feature - including one that is task-irrelevant - should update the shared perceptual state and therefore influence subsequent perception. In contrast, if history effects operate at the level of feature-specific readouts, updating should depend on whether the feature is selected for report, and thus the influence of an uncued feature should be minimal. To distinguish between these alternatives, we examined how judgments on trial *n* depend on the preceding stimulus when the reported feature switches between trials.

Figure 2C showed that when intensity was estimated on both trials, perceptual estimates were strongly attracted toward the previous percept. Figure 4A-B illustrates the corresponding case following a task switch, in which intensity was estimated on trial *n* but duration was estimated on trial *n-1*.

**Figure 4.**
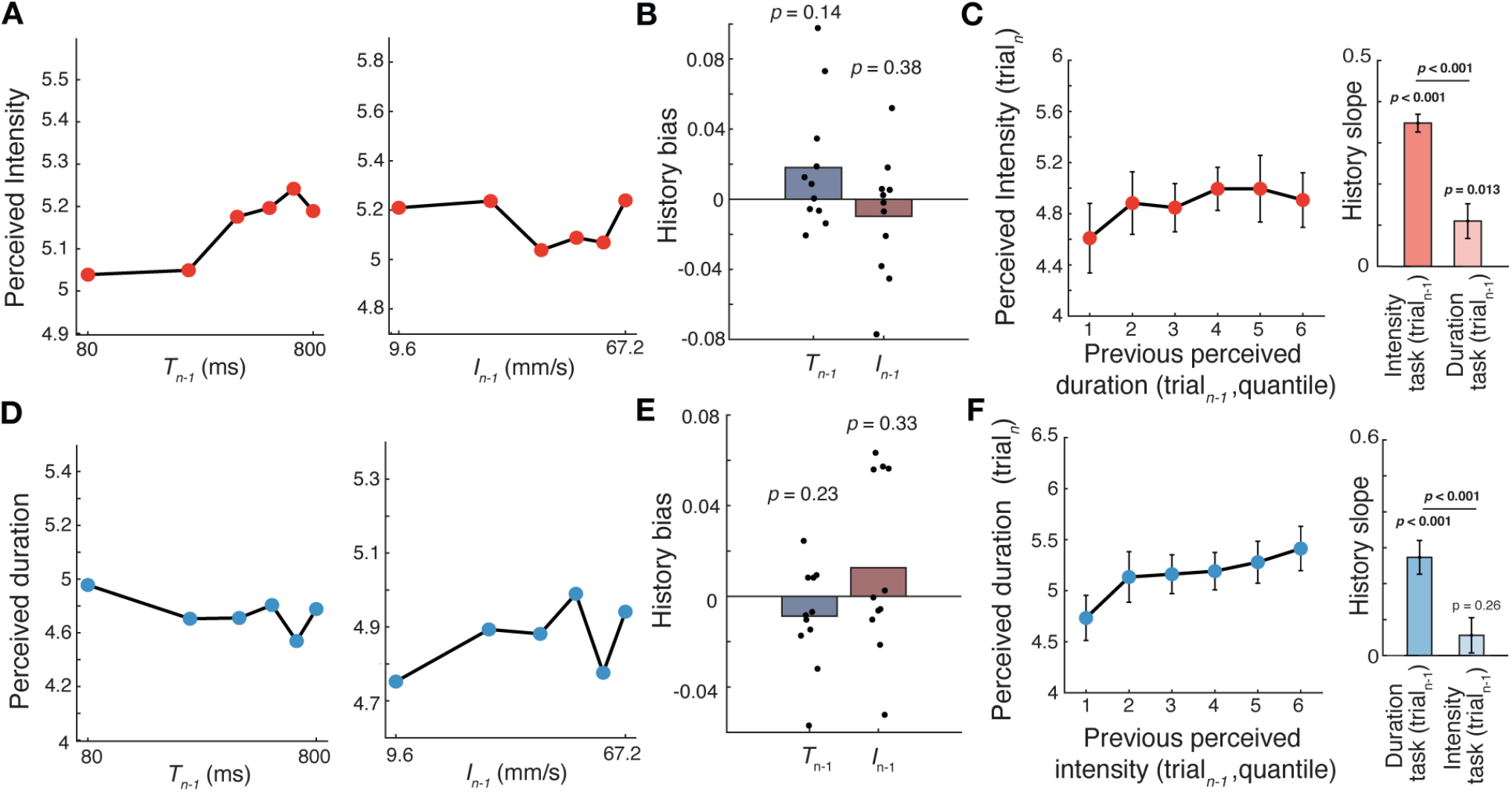
Serial dependencies when the cued feature is switched on consecutive trials. **(A)**Mean perceived intensity on trial *n* plotted against the duration (left) and intensity (right) of stimulus *n-1* when trial *n-1* required duration estimation. **(B)**Duration-history bias (blue) and intensity-history bias (red); each point is one subject. *p*-values are permutation tests against zero. Neither bias differs significantly from zero. **(C)**Left: Mean perceived intensity on trial *n* plotted against the perceived duration on trial *n-1*. Right: Slopes of previous-trial perceptual quantile versus current percept, for intensity task and duration task on trial *n-1*. Bar: average slope; error bar: S.E.M. **(D)**Mean perceived duration on trial *n* plotted against the duration (left) and intensity (right) of stimulus *n-1* when trial *n-1* required intensity estimation. **(E)**Duration-history bias (blue) and intensity-history bias (red); each point is one subject. *p*-values are permutation tests against zero. Neither bias differs significantly from zero. **(F)**Mean perceived duration on trial *n* plotted against the perceived intensity on trial *n-1*. Right: Slopes of previous-trial perceptual quantile versus current percept, for intensity task and duration task on trial *n-1*. Bar: average slope; error bar S.E.M.

In this condition, perceived intensity showed no detectable bias from either the duration or intensity of the preceding stimulus. When responses were expressed in perceptual coordinates, only a weak modulation by the previously perceived duration was observed (Figure 4C). Thus, attraction toward the preceding percept occurs when the cued feature is repeated (Figure 2C) but not following a task switch.

The same pattern holds when the task order is reversed. When trial *n-1* required intensity estimation and trial *n* required duration estimation, perceived duration on trial *n* was not significantly affected by either the intensity or duration of the preceding stimulus (Figure 4D-E), nor by the previously reported perceived intensity (Figure 4F).

Recent work has suggested that the influence of past experience is updated as new evidence is progressively incorporated ^11,14,16^. If so, there will be an influence of not only *n-1*, but also earlier trials. To test this, we examined the influence of previous trials (*n-1, n-2, n-3*) on judgments on trial *n* across trial sequences in which the task remained (by chance) constant (Figure 5A). Serial dependence bias was quantified using a generalized linear model in which the current perceptual estimate was predicted from the previous-trial perceptual estimate, separately for each lag (see Methods). The analysis reveals robust effects extending back to *n-2* and *n-3* for both duration and intensity, consistent with a continuously updated history trace whose influence decays systematically with trial distance.

**Figure 5.**
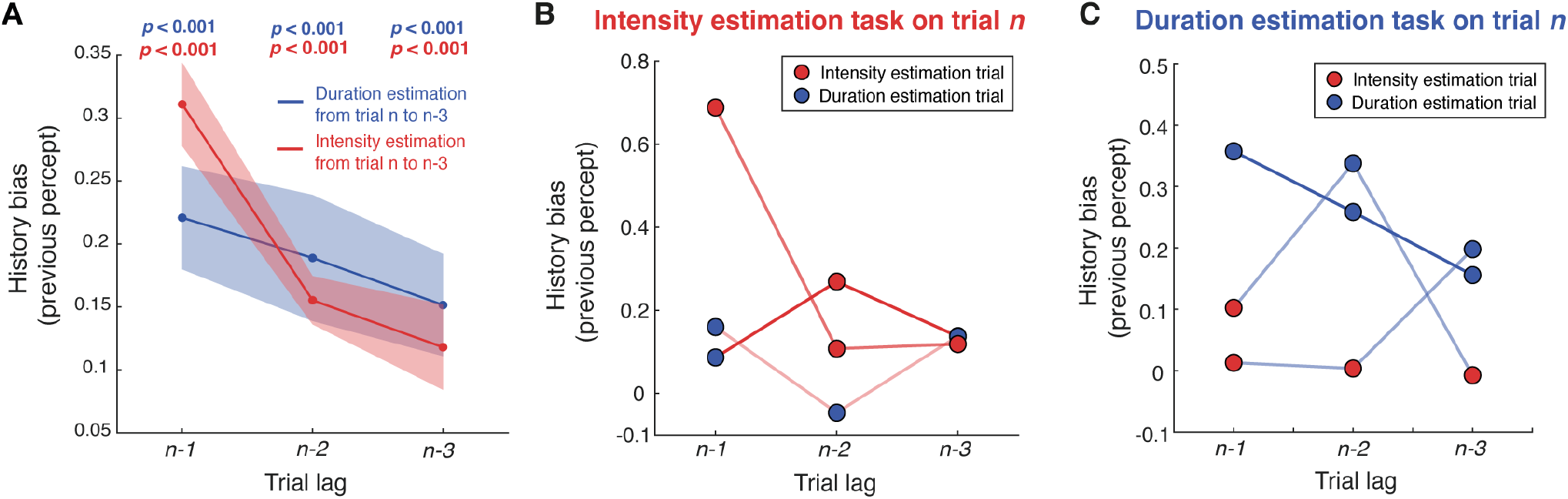
Serial dependencies across multiple preceding trials. **(A)** Bias on the perceived intensity (red) and perceived duration (blue) of trial *n* as a function of the perceived intensities and durations reported on trials *n-1, n-2*, and *n-3*. **(B–C)** Influence of perceptual reports on trials *n-1, n-2*, and *n-3* for different task-sequence combinations on the perceptual report of trial *n* intensity (B) and duration (C). Serial dependence was significantly modulated by task congruency for both intensity and duration judgments. Mixed-effects analyses revealed a significant Lag × Congruency interaction in both tasks (intensity: F(1,260)=26.87, *p* < 0.001; duration: F(1,260)=7.31, *p* = 0.007), indicating that the influence of past trials decreased across lags more strongly for congruent than incongruent task histories. (see Supplementary Figure 4 for all remaining patterns across trials)

Figure 4 indicates that the history trace of a given feature is updated only when that feature is extracted, as instructed by the cue. This implies that, across longer sequences, only those preceding trials matching trial *n* in cued task will affect trial *n* judgment. Figures 5B-C confirm this prediction for sequences comprising trials *n-1, n-2*, and *n-3*. In the intensity task (Figure 5B), perceived intensity on trial *n* showed little or no bias from perceived durations reported on trials *n-1* or *n-2*, yet remained reliably biased by previously reported intensities, including those on trials *n-2* and *n-3*. A parallel pattern is observed for duration estimation (Figure 5C): perceived duration on trial *n* is attracted toward previously perceived durations, but not toward intensities reported on intervening trials.

To quantify these effects, we fit linear mixed-effects models with history bias as the dependent variable and congruency, lag, and their interaction as fixed effects. In both tasks, history biases were significantly stronger following congruent than incongruent trials (intensity: F(1,26396) = 62.8, *p* < 0.001; duration: F(1,26179) = 15.03, *p* < 0.001 ). Critically, history biases decreased with lag, and this decay depended on task congruency, as reflected by significant Lag × Congruency interactions (intensity: F(1,26396) = 104, *p* < 0.001 ; duration: F(1,26179) = 124.9, *p* < 0.001). These results indicate that serial dependence is strongest for recent trials and is selectively enhanced when preceding and current judgments involve the same task dimension.

Together, these findings show that task-relevant history is not reset when an irrelevant feature is processed on intervening trials. Instead, feature-specific history traces for perceived duration and intensity persist and are selectively updated when each feature is probed, allowing one trace to remain stable across multiple trials even as the other is engaged. This demonstrates that intensity and duration histories are tracked in parallel and updated independently over time, consistent with distinct, feature-specific representations.

#### Computational model with feature-specific history traces

We extended previous formulations of history effects in working- and reference-memory tasks buffers^16,33,34^ to develop a computational model that (i) describes the transformation of sensory input into perceptual estimates, (ii) tracks their evolution into dynamic memory representations, and (iii) accounts for history-dependent biases in perceptual judgments. Within each perceptual dimension, two interacting components evolve continuously over time. Each newly formed percept is encoded in a Short-Term Buffer (STB), a transient trace representing the most recent percept and overwritten on each trial (Figure 6A). The STB interacts with a Long-Term Buffer (LTB), which encodes an accumulated representation of past percepts and serves as a perceptual prior. Over time, the STB is attracted toward the LTB, while the LTB is reciprocally updated by the current percept. Unlike the STB, the LTB is not reset but evolves continuously across trials, allowing successive stimuli to reshape the long-term representation and give rise to history-dependent biases.

**Figure 6.**
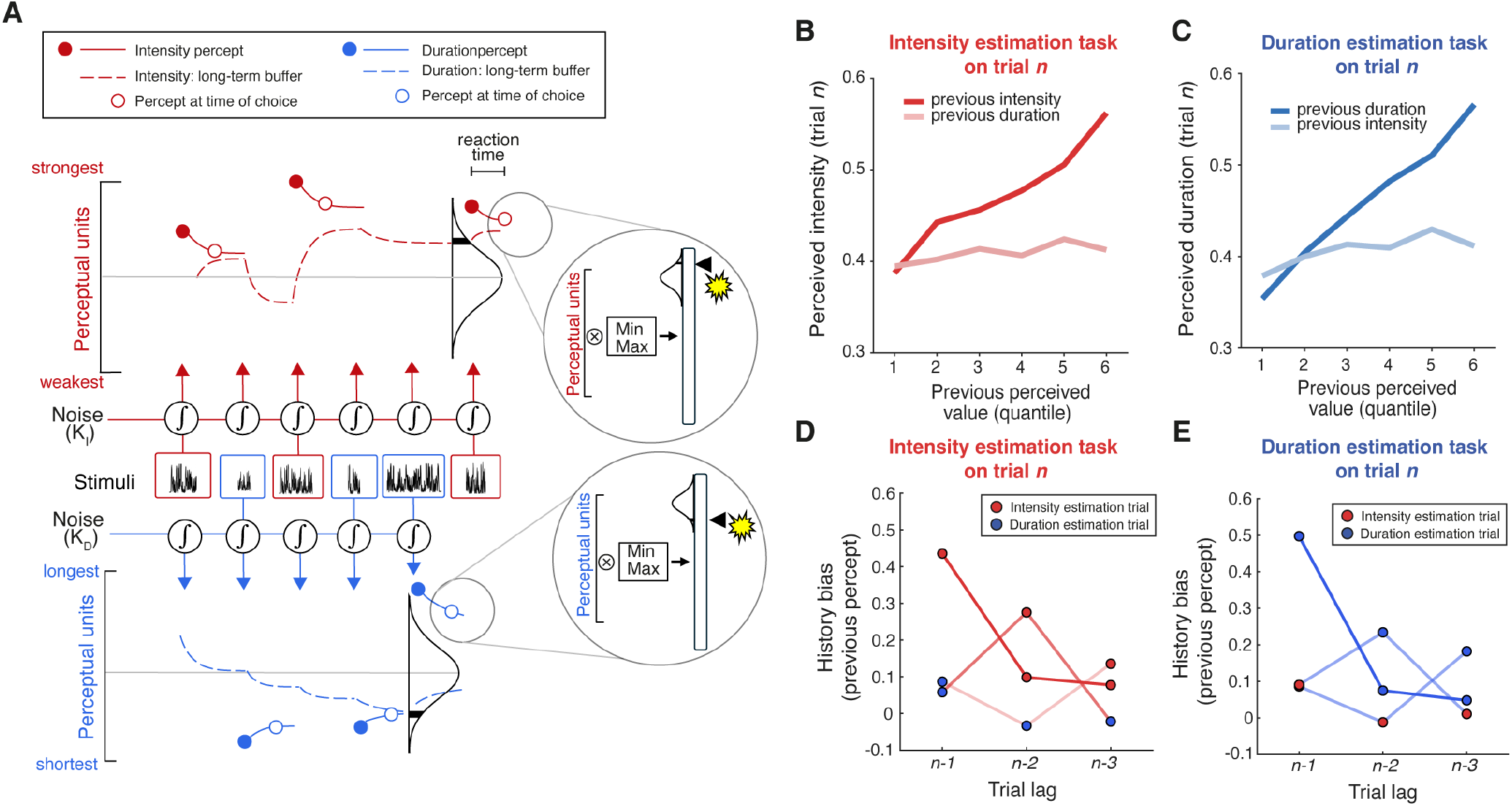
Computational model for history biases. **(A)** Six sequential tactile stimuli (intensity estimation, red; duration estimation, blue). Feature-specific history traces evolve in parallel: on intensity-cued trials, the intensity trace integrates the corresponding percept while the duration trace receives baseline drive (K_D_), and vice versa for duration-cued trials (the intensity trace receives baseline drive K_I_). Perceptual magnitudes are represented along vertical axes, with total accumulated history forming a Gaussian distribution (see ^16^). At stimulus onset, the instantaneous percept (filled circle) enters a short-term buffer, which is attracted toward the LTB while the LTB is reciprocally updated. The remembered value at response is shown as an open circle. Recent high intensity (top) and short duration (bottom) estimates shift the corresponding traces in opposite directions, simultaneously. and bias subsequent estimates. Behavioral reports (magnified inset) reflect a bounded readout of the internal estimate. **(B–C)** Model predictions of serial dependence: current estimates as a function of the previous-trial percept (quantile bins), showing attractive bias for the same feature. **(D–E)** Lag-dependent effects (*n-1* to *n-3*): the model predicts persistent, feature-specific traces that are not reset by intervening trials in which the other feature is reported.

#### Selective updating of parallel history traces

The experimental results place a constraint on how these history representations are updated. Although both perceptual dimensions are available on every trial, only the reported feature contributes to subsequent bias, indicating that perceptual history is not updated automatically from all available percepts. Instead, updating depends on whether a percept is actively accessed for judgment (Figures 4-5). To capture this constraint, the model implements parallel STB-LTB systems for each perceptual dimension, but restricts stimulus-driven updating to the task-relevant dimension. On each trial, the stimulus gives rise to a perceptual estimate, *P*_*n*_ which is encoded in the STB and incorporated into the corresponding LTB only when that feature is selected for report. In contrast, the non-relevant dimension evolves in the absence of stimulus-driven input, following baseline dynamics. This selective updating allows each history trace to persist across intervening trials without being overwritten by unattended percepts, accounting for the observed independence of intensity and duration history effects. The behavioral response reflects the STB value at the end of the post-stimulus delay (Figure 6A), mapped onto a bounded response scale via min–max normalization (see Methods).

At first glance, the observation of attraction to recent that are at the edges of the stimulus distribution (e.g., a tendency to judge “long” after a preceding “long” judgment) may seem at odds with the well-known phenomenon of contraction toward the mean^35^. This formulation reconciles the apparent contradiction: rather than being drawn toward a fixed global mean, the STB is attracted toward the current state of the LTB, which itself reflects recent perceptual history. Because the LTB evolves over time, contraction occurs toward a moving reference, giving rise to apparent attraction toward recent percepts.

#### Model behavior and correspondence with empirical data

Model parameters were optimized by minimizing the discrepancy between single-trial model predictions and observed behavioral responses using two-fold cross-validation (see Methods; Supplementary Figure 5). The model reproduces three central features of the behavioral data. First, the leaky integration stage generates interacting perceptual representations of intensity and duration (Supplementary Figure 6). Second, reciprocal STB–LTB interactions produce sequential dependencies: perceptual estimates on the current trial are attracted toward the long-term representation when the same perceptual dimension is reported on consecutive trials, whereas this influence is reduced following task switches (Figure 6B–C). Third, the model captures the key empirical finding: independent updating of intensity and duration priors across task sequences. Consistent with the results shown in Figure 5, the influence of percepts from *n-1, n-2*, and *n-3* depends on the task sequence but is not abolished by intervening trials in which the other feature is extracted. Instead, the corresponding history trace persists in the background and continues to evolve, shaping behavior when that feature is probed again (Figure 6D-E).

## Discussion

Perception of vibrotactile stimuli is shaped not only by current sensory input but also by recent history^11,14–16^. The aim of this study was to determine how history operates when multiple perceptual features of the same stimulus are extracted. Resolving this question is critical for understanding how the brain constructs and updates perceptual priors in multidimensional environments, and for constraining models of serial dependence.

When observers switched between judging the intensity and duration of a single vibrotactile stimulus (due to randomized task cue), perceived intensity was selectively attracted toward previously perceived intensities, and perceived duration toward previously perceived durations, with little cross-feature transfer. History effects were better predicted by previous perceptual reports than by previous stimulus features, and individual differences in the magnitude of the irrelevant feature’s influence in the current trial predicted the magnitude of the carry-over of the irrelevant feature to the next trial. Together, these results indicate that perceptual history is maintained in parallel, dimension-specific traces rather than as a single representation of recent stimuli.

An alternative account worth considering is that the task cue actively suppresses the irrelevant feature dimension, either prior to or during, perceptual processing. Under this view, feature-specific traces could in principle be updated whenever their associated feature is represented, but the task-dependent suppression of the unattended dimension in our paradigm prevents it from contributing to history. Such suppression may play a functional role in this task, where the physical features - duration and intensity - are encoded together, and their interaction can degrade perceptual accuracy. Distinguishing between these possibilities - readout-gated updating versus earlier suppression of task-irrelevant perceptual dimensions - will require paradigms in which both perceptual estimates are formed from the sensory input, but only one is selected for action.

Within a trial, intensity and duration influenced each other bidirectionally: stronger stimuli felt longer and longer stimuli felt stronger, replicating previous work in humans and rodents ^14,29,31,32,36^. This interaction suggests that both percepts rely on partially shared accumulation mechanisms^28,30^. Thus, while cross-feature interactions arise during current evidence integration, serial dependence reflects updating of feature-specific internal estimates. These findings show that perceptual continuity does not operate at the level of the stimulus object. Instead, temporal stability is imposed separately on the perceptual variables that are extracted.

The observation that history effects were anchored to previous percepts rather than to the physical properties of the stimulus is consistent with growing evidence that serial dependence reflects continuity in internal representations rather than persistence of sensory input^19,27,37^. Previous work has reported feature-specific serial dependence, with biases confined to the task-relevant dimension in domains such as numerosity and duration^22,38^. Building on this, our direct estimation paradigm reveals that separate perceptual history traces are updated in parallel and anchored to previous perceptual estimates rather than to physical stimulus properties.

Serial dependence is often interpreted in Bayesian terms, where recent percepts contribute to a prior that is combined with current sensory evidence^20,25^. Our findings support this framework but place an constraint on its structure. If intensity and duration relied on a common magnitude prior, as proposed by theories of a generalized magnitude system^39,40^, history effects should generalize across dimensions. Independent history traces for intensity and duration challenges accounts positing a unified magnitude prior and supports models in which separate priors are maintained for different perceptual variables^41^.

To understand how such feature-specific priors might arise, we developed a computational model in which perceptual estimates emerge from the interaction between two dynamical buffers operating on different time scales, a short-term buffer (STB) and a long-term buffer (LTB). After stimulus offset, the two buffers interact through bidirectional attraction: the STB gradually relaxes toward the LTB, while the LTB is updated by the newly formed percept. Through this interaction, the LTB tracks the recent distribution of perceptual estimates and effectively implements a history-dependent trace that biases subsequent percepts. Importantly, the model includes separate buffers for intensity and duration, leading to a gradual updating of feature-specific perceptual history. The model provides a mechanistic interpretation of Bayesian perceptual priors as dynamical states allowing slow accumulation of perceptual readouts and an ongoing interaction with current sensory estimates, rather than requiring explicit storage of previous stimuli or decisions.

More broadly, the locus of serial dependence remains debated. Several studies have reported signatures consistent with low-level sensory contributions, including stimulus-specific biases in early visual cortex and feature-selective carryover effects^24,42^. In contrast, other work shows that serial dependence depends on task relevance, attention, and working-memory delay, suggesting a post-perceptual origin at the level of memory or decision ^3,23,43^. Our findings help reconcile these views. History effects were not explained by the physical properties of previous stimuli but by previously reported perceptual estimates, indicating that temporal continuity operates on internal representations rather than on sensory input itself. At the same time, these representations remained selective for the task-relevant perceptual dimension, even when multiple percepts were extracted from the same physical event. Thus, serial dependence appears to act at the level of integrated perceptual variables – intermediate between early sensory encoding and decisional output. Some accounts have proposed that serial dependence reflects “continuity fields” that promote perceptual stability over time by integrating successive sensory inputs belonging to the same object or event^21^. Our results argue against a purely object-based form of continuity: even when intensity and duration were extracted from the same physical stimulus, their history effects remained dissociable and were updated independently according to task relevance.

Neurophysiological studies support this interpretation. In rodents performing vibrotactile judgments, neurons in posterior parietal cortex encode stimulus history and causally contribute to contraction bias. In primates and humans, serial dependence has been linked to activity patterns in parietal and prefrontal cortex, consistent with working-memory-like population dynamics^44^. Intensity and duration percepts are themselves thought to arise from temporal integration processes with distinct time constants in higher-order somatosensory–parietal networks^14,28,30^. Consistent with this interpretation, recent recordings in rats performing a tactile intensity reference-memory task showed that frontal cortical populations jointly encode current sensory inputs and prior sensory context, providing a potential neural substrate for the history-dependent perceptual states captured by our model^45^. Together, these findings suggest that perceptual history reflects recurrent dynamics in higher cortical circuits that maintain feature-specific decision variables. Recent work in zebrafish identified a thalamus–brainstem attractor network underlying history-biased decisions^46^. While this architecture resembles the interacting buffers proposed here, our findings suggest that in humans the maintained state reflects task-relevant percepts rather than physical stimulus features, leaving open whether the relevant history traces are stored in subcortical circuits or in cortical networks encoding perceptual variables.

Recurrent network models provide a mechanistic account of how perceptual priors emerge from ongoing neural dynamics. Slow decay, short-term plasticity, or adaptive gain can produce attractive biases toward recent states without explicit storage^44,47^. Recent work shows that recurrent neural networks trained in temporally structured environments naturally develop history-dependent biases^48^. Our results suggest that such mechanisms must operate in parallel across partially segregated circuits, enabling independent updating of multiple perceptual priors from a common sensory input.

Altered use of recent sensory history has been reported in several clinical conditions such as schizophrenia^18^, psychosis^49^ and autism^50^, possibly due to NMDA-R disfunctions^7,18^. These findings suggest that symptoms may reflect abnormalities in how perceptual memory traces are constructed. Our results may help constrain models of perceptual history formation which could provide new insights into the mechanisms underlying alterations of perceptual history biases in clinical conditions.

## Methods

### Participants

#### Ethics Statement

Human subjects were tested after giving their written informed consent. Protocols conformed to international norms and were approved by the Ethics Committee of SISSA (protocol number 6948-II/7).

#### Human subjects

Eleven healthy human subjects (age range 22–35 yrs) were for the direct estimation tasks. All subjects were recruited on-line through the SISSA Sona System. (https://sissa-cns.sona-systems.com/).

### Stimulus and behavioral task

#### Stimulus generation

Vibrations were generated by stringing together sequential velocity values (*v*_*t*_) at 10,000 samples/s, taken from a normal distribution. Converting *v*_*t*_ to its absolute value, *sp*_*t*_, the distribution takes the form of a folded, half-Gaussian. A Butterworth filter with 150 Hz cutoff was then applied to yield low-pass filtered noise. The *sp*_*t*_ time series for a given trial was taken randomly from among 50 unique sequences of pseudo-random values. Because stimuli were built by sampling a normal distribution, the statistical properties of an individual vibration did not perfectly replicate those of the distribution from which it was constructed. As a vibration’s actual mean speed could not precisely match the distribution from which it was sampled, the assigned value may be considered “nominal” mean speed, referred to as *I*.

#### Direct estimation task

Each subject went through two separate session, held on different days. Each session began with a training phase. In this phase, subjects received 40 stimuli, sampled randomly from the 100 possible stimuli (10 possible *I* values from 9.6 mm/s to 67.2 mm/s and 10 possible *T* values from 80 to 800 ms, linearly spaced), to become confident with the task and to sample the stimulus range. In the test phase, a single stimulus was presented, characterized by intensity, *I*, and duration, *T*. 500 ms before each stimulus presentation a visual cue, either a blue or a red square, was presented. either before or after the delivery of the vibration. The color of the visual cue instructed to report either the perceived duration (blue) or perceived intensity (red) of the presented stimulus. After a post-stimulus delay of 500 ms, a slider appeared on the screen. The slider did not present any landmark, ticks or numbers. The orientation of the slider was randomly changed at each trial, either vertical or horizontal. Subjects were instructed to report the perceived intensity or else the perceived duration of the vibration on a subjective scale, in which the extreme left/bottom position indicated a very weak or a very short stimulus, and the extreme right/top position indicated a very strong or a very long stimulus. The report was done by mouse-clicking on the chosen position along the slider. A total of 800 trials was presented at each session.

### Behavioral analyses

#### Normalization of direct estimation task data

Results from the direct estimation task were analyzed taking into account that not all subjects used the whole range of the slider; every participant set their minimum and maximum responses at a different position in the scale. In order to make each subject’s subjective scale comparable, we used a min-max normalization algorithm:

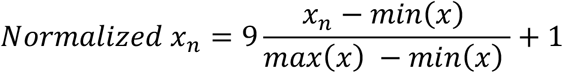

where *x*_*n*_ is the non-normalized response on trial *n, x* is the range of total responses, and *Normalized x*_*n*_ is the normalized response. We then multiplied by 9 and added 1, so that the normalized responses range from 1 to 10.

#### Quantification of serial dependencies between consecutive trials

To quantify serial effects across trials, we examined how the reported percept on trial *n* depended on the stimulus presented on the immediately preceding trial (*n* − 1). For each task, average perceptual reports *y* were computed as a function of the previous trial’s stimulus intensity *I*_*n-1*_ and duration *T*_*n-1*_. Specifically, for a given stimulus dimension *x*,

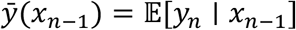

where *x* corresponded either to stimulus duration *T* or vibration intensity *I*. To quantify the magnitude of history dependence at the individual-subject level, log–log linear regressions were fitted between the previous-trial stimulus magnitude and the current perceptual report:

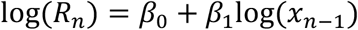

The regression slope *β*_1_ was used as a scalar measure of perceptual history bias. Separate slopes were estimated for previous duration (duration history biases) and previous intensity (intensity history biases) effects in each task. The analysis was ran separately for trials in which the cued feature is repeated (Figure 2) or switched (Figure 4) on consecutive trials.

#### Comparison between physical-history and perceptual-history models

To determine whether sequential biases were better explained by the physical properties of the previous stimulus or by the previous perceptual estimate, we compared two generalized linear models using cross-validated prediction accuracy. Analyses were performed separately for the intensity and duration tasks. Only consecutive trials from the same task were included. Behavioral responses were normalized within session and task and log-transformed before modeling. For each participant, two competing models were fitted. The physical-history model predicted the current response *y*_!_ from the current relevant stimulus dimension, the current irrelevant dimension, and both relevant and irrelevant stimulus features from the previous trial:

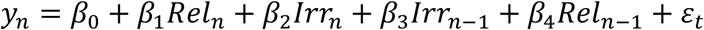

where *Rel* denotes the task-relevant stimulus dimension (intensity in the intensity task; duration in the duration task), and *Irr* denotes the task-irrelevant dimension.

The perceptual-history model replaced previous physical stimulus terms with the observer’s previous perceptual report:

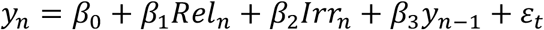

All predictors and responses were z-scored within the training folds prior to fitting. Models were fitted using Gaussian generalized linear regression (glmfit, MATLAB). Predictive performance was evaluated using 5-fold cross-validation. For each fold, models were trained on *K* − 1 partitions and tested on the held-out partition. Predictions were transformed back to the original log-response space before computing cross-validated explained variance:

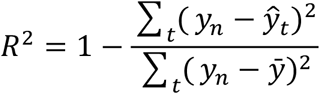

Model comparison was quantified as:

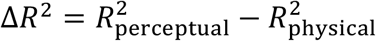

Positive Δ*R*^2^ values indicate superior predictive performance of the perceptual-history model.

#### Relationship Between Sensory and History Biases

To quantify the relationship between sensory interference and sequential-history effects, we fit a linear mixed-effects regression model across all trials:

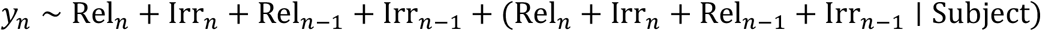

where *y* denotes the reported percept, Rel_*n*_ and Irr_*n*_ correspond to the current relevant and irrelevant stimulus dimensions, and Rel_*n*-1_ and Irr_*n*-1_ denote the corresponding features from the previous trial. Subject-specific random slopes associated with current irrelevant features (Irr_!_) and previous irrelevant features (Irr_*n*-1_) were then extracted and correlated across participants to assess whether individuals showing stronger cross-dimensional sensory interactions also exhibited stronger cross-dimensional history biases.

### Lag-dependent history analysis

To characterize the temporal structure of sequential biases, we quantified the influence of perceptual reports from the previous three trials (*n* − 1, *n* − 2, *n* − 3) separately for the intensity and duration tasks. Previous trial sequences were grouped according to binary task-history patterns indicating whether each preceding trial belonged to the current task or the alternate task. For each lag, a value of 1 denoted a previous trial from the same task, whereas 0 denoted a trial from the alternate task. Selected task-history patterns were analyzed independently. First, for each participant and lag, we estimated history bias using separate linear regressions in which the current perceptual report was predicted by the perceptual report at a single previous lag while controlling for current stimulus duration and intensity:

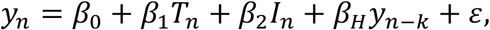

where *y*_*n*_ denotes the current perceptual report, *T*_*n*_ and *I*_*n*_ denote the current stimulus duration and intensity, respectively, and *y*_*n−k*_ is the perceptual report from lag *k*(*k* = 1,2,3). All predictors were z-scored within participant prior to fitting. Later, for each participant and pattern, current perceptual reports were modeled using multiple linear regression in which the current perceptual report was predicted by the perceptual report at all three previous lags while controlling for current stimulus duration and intensity:

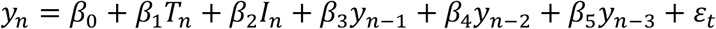

where *y*_*n*_ denotes the current perceptual estimate, *T*_*n*_ and *I*_*n*_ correspond to current stimulus duration and intensity, respectively, and *y*_*n−k*_ denotes the perceptual response from lag *k*. Current stimulus predictors and lagged perceptual responses were z-scored prior to regression.

For each participant, regression coefficients associated with the three lagged perceptual terms (*β*_3_, *β*_4_, *β*_5_) were extracted as measures of history dependence across temporal lags.

To determine how serial dependence varied across temporal lags and task congruency, each lag was labeled as congruent or incongruent depending on whether the corresponding preceding trial involved the same task as the current trial. History-bias coefficients were analyzed using a linear mixed-effects model with fixed effects of lag, congruency, and their interaction, random intercepts and lag slopes for subjects, and random intercepts for congruency-pattern sequence:

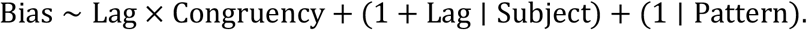

Statistical significance of fixed effects was evaluated from the mixed-effects model.

### Behavioral model

#### Model overview and fitting procedure

Participants completed two experimental sessions. We modeled behavior using two complementary components: a perceptual model and a history model. The perceptual model was adapted from Toso et al^28^ described how stimulus intensity and duration were transformed into trial-wise latent perceptual representations through leaky temporal integration. The history model was adapted from^33,34^ and described slower sequential fluctuations in perception arising from interactions between short-term and long-term buffers (STB/LTB).

The models were fit hierarchically using two-fold cross-validation across sessions. First, the perceptual model was fit on one session to estimate the latent perceptual representations associated with each trial. These estimated perceptual states were then used as inputs to the history model, which was fit on the other session. This hierarchical approach dissociated stimulus-driven percept formation from slower history-dependent dynamics. The history model therefore operated on latent perceptual estimates generated by the perceptual model, allowing sequential dependencies to emerge from internal perceptual representations.

#### Perceptual model architecture

On each trial *n*, a vibrotactile stimulus is characterized by its physical intensity *I*_*n*_ and duration *T*_*n*_. The vibration signal is modeled as a sequence of instantaneous velocity samples drawn from a half-Gaussian distribution with mean *sp*_*n*_. Perceptual estimates are generated through a leaky integration process applied to this sensory drive *u*_*n*_,

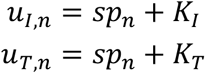

where *K*_1_, *K*_*D*_ captures task-specific baseline accumulation in the absence of stimulus input. The temporal evolution of the internal percept follows Ornstein–Uhlenbeck dynamics, yielding at stimulus offset:

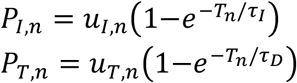

where *τ*_*I*_ and *τ*_*D*_ are temporal integration constants controlling the accumulation of sensory evidence over time. Critically, the attended dimension receives stimulus-dependent input, whereas the unattended dimension evolves solely under baseline drive. Thus, during intensity-report trials, *P*_*I*_ reflects stimulus magnitude while *P*_*T*_ evolves from baseline activity alone; the converse holds during duration-report trials. Predicted perceptual values were normalized using a min-max procedure separately within each session and task before history model fitting.

#### History model architecture

The history model captured sequential dependencies through interactions between a short-term buffer (STB) and a long-term buffer (LTB), implemented independently for intensity and duration. Trial-by-trial perceptual estimates generated by the perceptual model (*P*_*I*_, *P*_*T*_) served as sensory inputs to the history system.

For each trial, buffer dynamics evolved across three phases: stimulus presentation, a post-stimulus delay (0.5 s), and a post-response/inter-trial delay (0.5 s). During the stimulus phase, the perceptual estimate was encoded into the relevant STB. Following stimulus offset, STB and LTB continued interacting during the post-stimulus delay, and the perceptual report was read out from the relevant STB state at the end of this delay period. The buffers then continued evolving during the inter-trial interval, generating carry-over effects onto subsequent trials.

The unattended dimension evolved according to the same STB/LTB dynamics as the attended dimension. However, because sensory evidence was absent in the unattended channel, its perceptual drive was determined solely by the baseline accumulation term (*k*_*I*_ or *k*_*T*_). Thus, unattended buffers continued updating across trials, but without task-relevant sensory input. STB and LTB dynamics followed coupled exponential interactions:

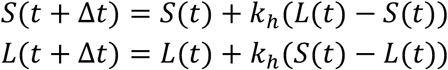

where *S* and *L* denote STB and LTB states, respectively. The interaction kernel was defined as:

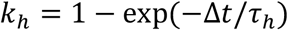

with separate history timescales fitted for intensity and duration (*τ*_*h*_,_*I*_, *τ*_*h*_,_*D*_).

#### Optimization

Model parameters were estimated by minimizing the negative log-likelihood of behavioral reports using constrained nonlinear optimization (fmincon, MATLAB). Optimization was repeated across 1000 random restarts, and the parameter set yielding the lowest negative log-likelihood was retained. For the perceptual model, free parameters included the perceptual integration timescales (*τ*_*I*_, *τ*_*T*_) and baseline accumulation terms (*k*_*I*_, *k*_*T*_). Timescale parameters were optimized in log-space to enforce positivity, with bounds:

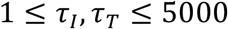

and baseline accumulation parameters constrained to:

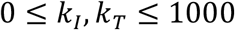

Initial conditions were randomly jittered around predefined starting values for each restart. For the history model, separate history timescales were fitted for intensity and duration (*τ*_*h,I*_, *τ*_*h,T*_). These parameters were also optimized in log-space with bounds:

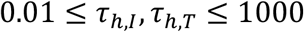

## Funding

Human Frontier Science Program, project RGP0017/2021 (MD)

Italian Ministry PRIN 2022 contract 20224FWF2J (MD)

Italian Ministry PNRR contract P20229752W (MD)

Regional Laboratory for Advanced Mechatronics, LAMA FVG (http://lamafvg.it)

European Union Horizon 2020 MSCA Programme grant 813713 (MED)

## Author contributions

Conceptualization: AT, MED; Methodology: AT, MED; Investigation: AT; Visualization: AT, MED Funding acquisition: MED; Project administration: MED; Supervision: MED; Writing – original draft: AT, MED; Writing – review & editing: AT, MED

## Competing interests

None

## Data and code availability

Custom-written code and data are available at: https://github.com/Tosot91/Independent-history-traces-for-distinct-percepts-derived-from-a-single-stimulus.git

## Notes

### Competing Interest Statement

The authors have declared no competing interest.

