## Supplementary information for "Independent history traces for distinct percepts derived from a single stimulus"

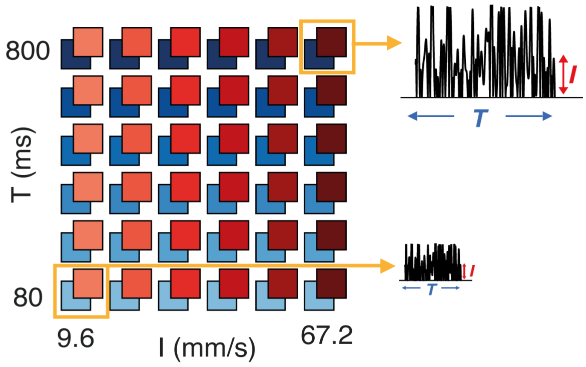


**Supplementary Figure 1. Direct estimation task** Stimulus matrix for direct estimation task. The vibration duration and intensity was randomly picked from the set of ($T$, $I$) combinations represented by the colored squares. Two sample stimuli from the upper right and lower left of the matrix are illustrated.

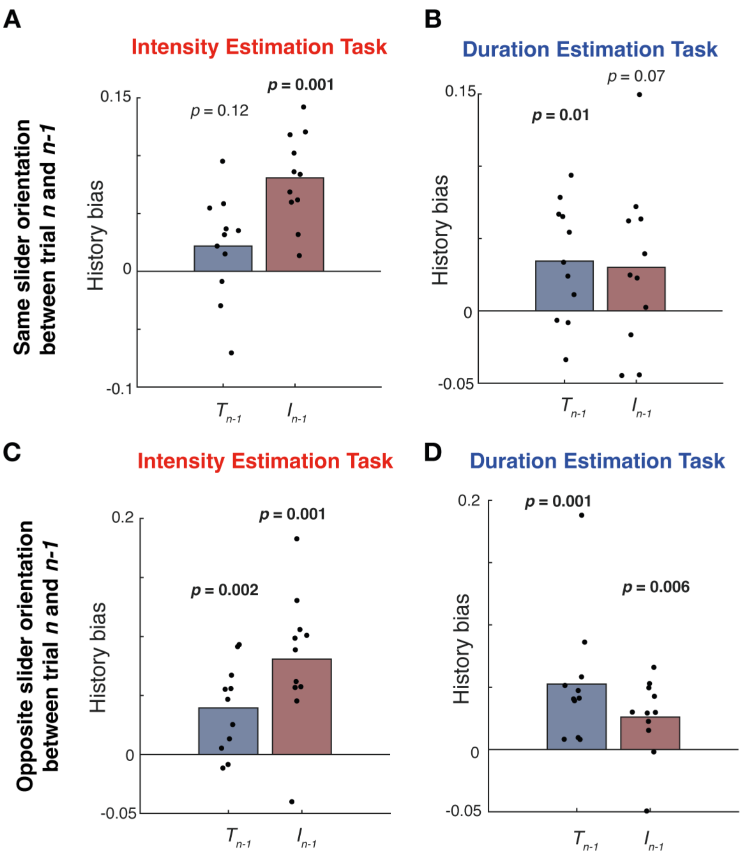


**Supplementary figure 2. Serial dependencies in the direct estimation task for different slider orientations. (A-B)** Trials in which the same slider orientation between trial *N* and trial *N-1* was presented: (A) shows the *duration history bias* in blue and *intensity history bias* in red of all subjects for intensity estimation trials. Each point is a single subject. (B) panel shows symmetrical effects for duration estimation trials. **(C-D)** Trials in which opposite slider orientations between trial *N* and trial *N-1* were presented: (C) shows the *duration history bias* in blue and *intensity history bias* in red of all subjects for intensity estimation trials. Each point is a single subject. (D) shows symmetrical effects for duration estimation trials.

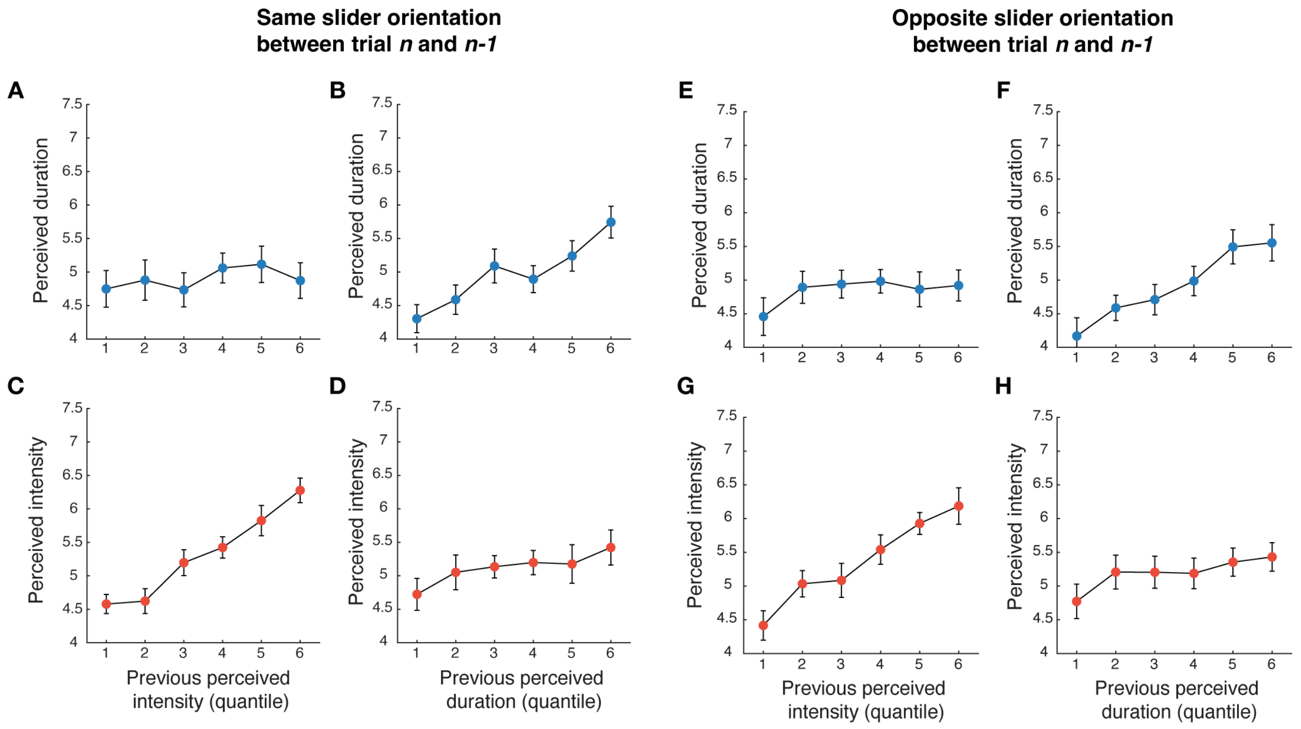


**Supplementary figure 3 Serial dependencies in the direct estimation task: different tasks on current and previous trial, for different slider orientation** (A-B) Duration estimation trials in which the same slider orientation between trial *N* and trial *N-1* was presented: mean perceived duration reported at trial *N* as a function of perceived intensity (A) and perceived duration (B) of the stimulus presented on Trial *N-1*. **(C-D)** Intensity estimation trials in which the same slider orientation between trial *N* and trial *N-1* was presented : mean perceived intensity reported at trial *N* as a function of perceived intensity (C) and perceived duration (D) of the stimulus presented on Trial *N*. **(E-H)** same as (A-D) for trials in which opposite slider orientations were presented between trial *N* and trial *N-1*.

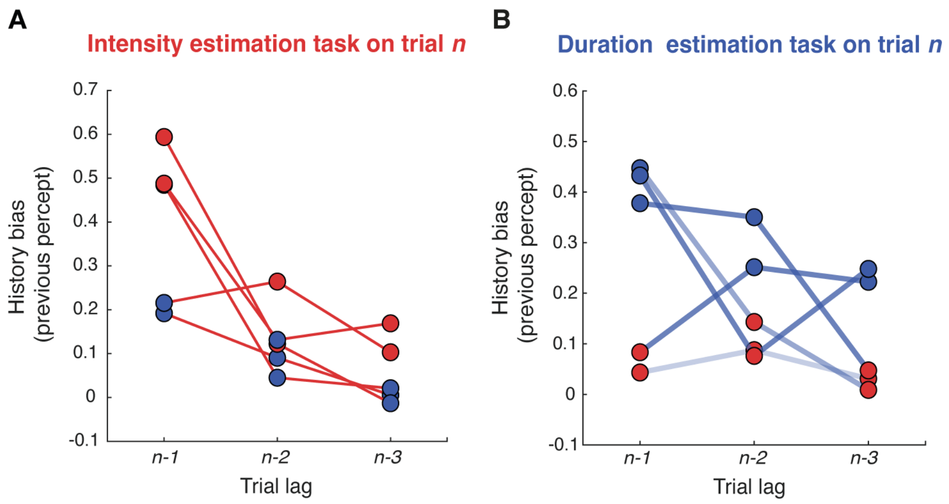


**Supplementary figure 4. Serial dependencies across multiple preceding trials (A-B)** Influence of the percepts reported on trials *n-1*, *n-2*, and *n-3* – for different combinations of task sequences (duration estimation trials are marked with blue points; intensity estimation in trials are marked with red points) – on the perceived intensity (panel A) or perceived duration (panel B) reported on trial *n*.

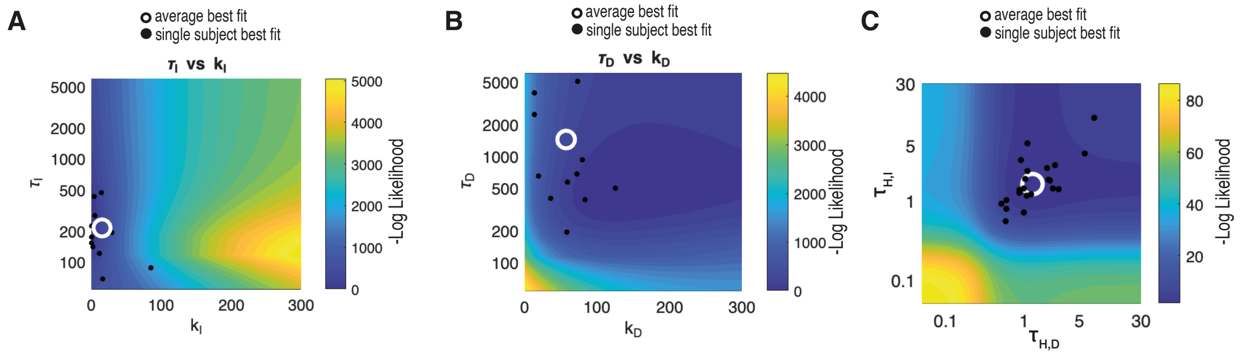


**Supplementary figure 5. Best fit parameters**. **(A-B)** Negative Log Likelihood for the perceptual model for different values of tau of integration and noise value K, for intensity (A) and duration (B) estimation trials. **(C)** Negative Log Likelihood for the history model for different values of tau of STB-LTB attraction for intensity and duration buffers. White circle indicates the average bets parameter fit for all subjects, individual black points indicate single subject parameters.

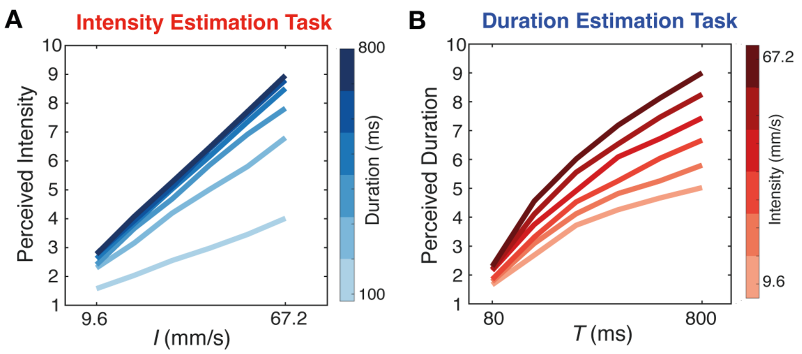


**Supplementary figure 6. Interaction between intensity and duration for model simulations.** Bias imposed by the irrelevant feature is captured by the computational model. Mean predicted perceived intensity (A) and perceived duration (B) increased a function of both stimulus intensity $I$ and duration $T$.
